# Deep learning to decompose macromolecules into independent Markovian domains

**DOI:** 10.1101/2022.03.30.486366

**Authors:** Andreas Mardt, Tim Hempel, Cecilia Clementi, Frank Noé

## Abstract

The increasing interest in modeling the dynamics of ever larger proteins has revealed a fundamental problem with models that describe the molecular system as being in a global configuration state. This notion limits our ability to gather sufficient statistics of state probabilities or state-to-state transitions because for large molecular systems the number of metastable states grows exponentially with size. In this manuscript, we approach this challenge by introducing a method that combines our recent progress on independent Markov decomposition (IMD) with VAMPnets, a deep learning approach to Markov modeling. We establish a training objective that quantifies how well a given decomposition of the molecular system into independent subdomains with Markovian dynamics approximates the overall dynamics. By constructing an end-to-end learning framework, the decomposition into such subdomains and their individual Markov state models are simultaneously learned, providing a data-efficient and easily interpretable summary of the complex system dynamics. While learning the dynamical coupling between Markovian subdomains is still an open issue, the present results are a significant step towards learning “Ising models” of large molecular complexes from simulation data.

---

The understanding of protein function is often interlinked with understanding protein dynamics. Molecular dynamics (MD) simulations are a valuable tool to study these dynamics on an atomistic level [1–6]. However, further methods are necessary to extract the statistically relevant information and to help overcome the discrepancy between feasible simulation length and the timescales of relevant processes. A common approach to enhance sampling of a specific process of interest is to bias the simulation along a reaction coordinate aligning with the process [7–13]. In comparison, the Markov modeling approach [14–20] extracts kinetic information and tackles the sampling problem without requiring the definition of few predefined reaction coordinates by combining arbitrary numbers of short unbiased distributed simulations to model the long-timescale behavior of target systems. Consequently, multiple software packages [21, 22] have been developed over the last decade providing assistance in estimating these models. They often include a pipeline for feature selection [21–24], dimension reduction [25–31], clustering [32–35], transition matrix estimation [15, 19, 36, 37], and coarse graining [38–44]. Markov state models (MSMs) have been applied to a wide range of molecular biology problems such as protein aggregation [45–47] or ligand binding [48–50] and can be a valuable tool to understand experimental data on the atomistic scale [51, 52].

The necessity to assess a model’s performance and thereby rank its quality encouraged the development of variational methods [53, 54], in particular the variational approach for Markov processes (VAMP) [55]. This variational formulation has allowed us to replace the aforementioned pipeline with an end-to-end deep learning framework called VAMPnet [56], which simultaneously learns a dimension reduction of the molecular system to the collective variables best describing the rare event processes and an MSM on these variables. The framework can be used to further drive MD simulations along these learned collective variables [57, 58]. We can also use this framework to estimate statistically reversible MSMs and incorporate constraints from experimental observables [59–61].

Despite these developments, there is a fundamental scaling problem in describing MD in terms of transitions between global system states: While the assignment of MD configurations to discrete global states representing the metastable groups of structures is an excellent model for small cooperative molecular systems, such as small to medium proteins, larger molecular systems (e.g. proteins with hundreds of amino acids) have an increasing number of subsystems whose dynamics are (nearly) independent [62] (Fig. 1). Consider, for example, a solution of *N* proteins which undergo transitions between their open and closed states independently when these proteins are dissociated and these transitions only (partially) couple when they are associated with other proteins. The number of global system states is 2^*N*^, i.e. grows exponentially with the number of subsystems N [63, 64]. This means any form of simulation or analysis which explicitly distinguishes global system states will not scale to large molecular systems.

**Figure 1:**
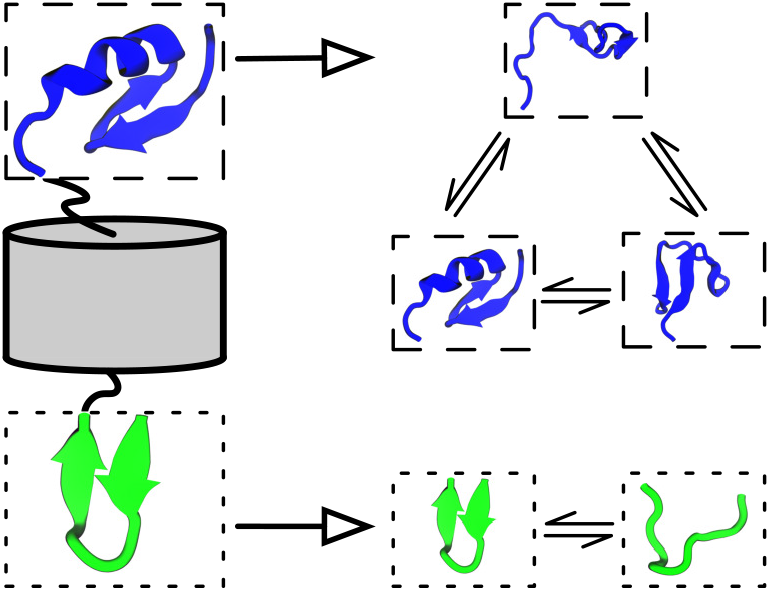
The iVAMP concept as visualized by modeling dynamics of a protein that has two independent, flexible regions separated by a rigid barrel. iVAMPnets learn an assignment of the C-(blue/top) and N-termini (green/bottom) into independent subsystems from molecular dynamics trajectories (left column). Moreover, the dynamics of both termini are modeled with statistically independent VAMPnets (right column).

At the same time, the (approximate) independence between subsystems is also key to the solution of the problem. A scalable solution needs to address two separate issues: (a) dividing the protein system into approximately Markovian subsystems and (b) learning the coupling between them. Olsson & Noé [63] made a first attempt at (b), by learning a dynamic graphical model between predefined subsystems. This approach leads to a graphical model, or Markov random field, resembling Ising or Potts models in physics, with the key difference that both the definition of the individual subsystems or “spins” as well as their transition dynamics need to be learned. In contrast, Hempel et al. [64] proposed a solution for (a) by approximating the global system dynamics as a set of independent (uncoupled) Markov models (termed Independent Markov decomposition, IMD). They furthermore propose a pairwise independence score of features, which allows to detect nearly uncoupled regions where independent Markov state models can be estimated subsequently.

In this manuscript, we present a joint IMD and VAMP approach (termed independent VAMPnet, or shorthand iVAMPnet) that significantly advances our ability to identify approximately independent Markovian subsystems (issue a) by generalizing IMD to neural network basis functions. iVAMPnets are an integrated end-to-end learning approach that decomposes the macromolecular structure into subsystems that are dynamically weakly coupled, and estimates a VAMPnet for each of these subsystems to promote a comprehensible analysis of the subsystem dynamics (Fig. 1). In comparison to previous implementations of IMD, our approach learns an optimal decomposition into independent subsystems and can find collective variables that are nonlinear combinations of the input features.

## Theory

### Markov state models and Koopman models

Markovian dynamics can be modeled by the transition density:

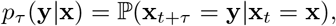

which is the probability density to observe configuration **y** at time *t* + *τ* given that the system was in configuration **x** at time *t*. Based on the transition density we can characterize the time evolution of a probability density *x* as:

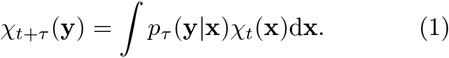

By discretizing the molecular state space in a suitable way and defining a transition matrix **T** between discrete states, we can linearize this equation as:

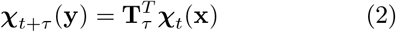

This is the equation of a Markov state model, where the element *i* of the vector ***χ***_*t*+*τ*_(**y**) is the probability to be in the discrete state *i* at time *t*+*τ*. Furthermore, the transition matrix elements (**T**_*τ*_)_*ij*_ describe the transition probabilities for jumping to state *j* given state *i* within a time *τ*. In the case of fuzzy state assignments, e.g., as with VAMPnets, Eq. (2) describes the more general Koopman model [65] and **T**_*τ*_ becomes the Koopman matrix. This means that probability densities are still propagated but the matrix elements cannot be interpreted as transition probabilities.

The lag time *τ* is common to all Markovian models and is usually chosen with the aid of an implied timescales test [66]. If a too small *τ* is chosen, the resulting model is not a valid Markov model (resulting in errors of the predicted variables) — a too large lag time produces a model that unnecessarily discards kinetic information. We therefore usually choose the smallest lag time above which the implied timescales are approximately constant.

We now seek to find a state assignment ***χ*** and model matrix **T** that satisfy Eq. (2) and also succeed in predicting the long-time behavior, i.e., for multiples of the lag time *τ*. Formally, ***χ*** are (initially unknown) basis functions, i.e., we assume that the relevant dynamic features can be expressed by a linear combination of them. VAMP [55] tells us that an optimal solution is reached when ***χ*** can span the left (*ψ*_1_,…, *ψ_k_*)^*T*^ and right singular functions (*ϕ_i_*,…, *ϕ_k_*)^*τ*^ of the transition operator. They can be found by maximizing the singular values of a matrix that can be estimated from simulation data (see Eqs. 8–12 in Methods). In the case of a VAMPnet [56], deep neural networks are trained by maximizing the VAMP score, so as to represent optimal fuzzy state assignments.

In equilibrium, the singular functions correspond to the eigenfunctions of the Markov state model and the singular values to its eigenvalues. As the Koopman model still propagates densities, it is instructive to inspect the eigenfunctions and implied timescales of **T** since they describe the slow dynamics of a given system.

### iVAMPnets and iVAMP-score

To implement iVAMPnets, we need to bridge the gap between the deep neural networks of VAMPnets and the spatial decomposition of independent Markov models. The general idea is to set up multiple parallel VAMPnets, each modeling the Markovian dynamics of a separate, independent subsystem of the molecule, together with an attention mechanism that identines these subsystems. Thus, each independent VAMPnet should only receive the time dependent molecular geometry features representing its specific subsystem. For example, such an attention mechanism could separate different protein domains and channel the data of individual domains to separate VAMPnets. We therefore develop an architecture that combines a meaningful attention mechanism and parallel VAMPnets and trains them with a loss function that simultaneously promotes dynamic independence between the subsystems and slow kinetics within each subsystem (Fig. 2). iVAMPnets are designed to optimize both these objectives simultaneously.

**Figure 2:**
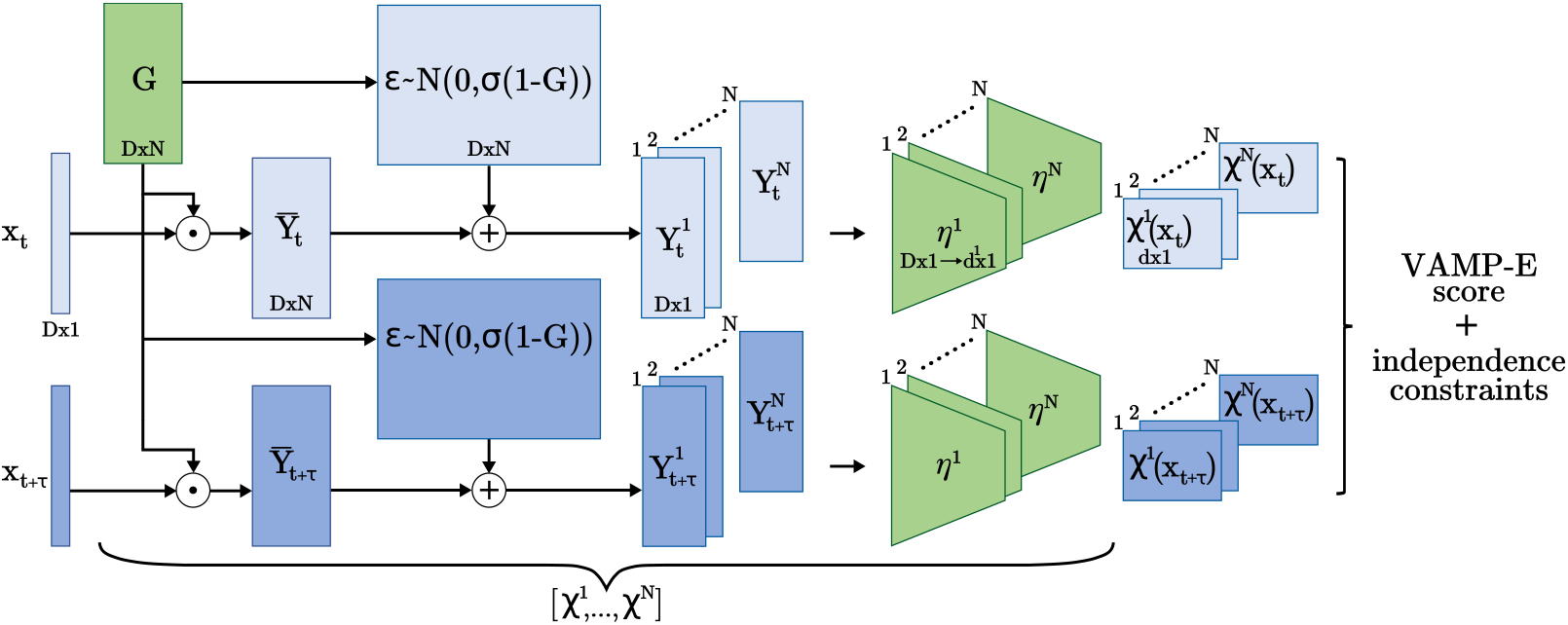
Architecture of an iVAMPnet for *N* subsystems, where trainable parts are shaded green. Two lobes are given for configuration pairs **x**_*t*_ and ***x***_*t*+*τ*_, where the weights are shared. Firstly, the input features are element wise weighted 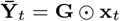 with a mask **G** ∈ ℝ^*D*×*N*^, where each subsystem learns its individual weighting. The mask values can be interpreted as probabilities to which subsystem the input feature belongs. In order to prevent the subsequent neural network to reverse the effects of the mask, we draw for each input feature *i* and subsystem *j* an independent, normally distributed random variable 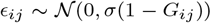. This noise is added to the weighted features 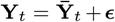. Thereby, the attention weight linearly interpolates between input feature and Gaussian noise, i.e., if the attention weight *G_ij_* = 1, *Y_ij_* carries exclusively the input feature *x_i_*, if *G_ij_* = 0, *Y_ij_* is simple Gaussian noise. Afterwards, the transformed feature vector is split for each individual subsystem 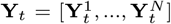 and passed through the subsystem specific neural network ***η***^*i*^. We call the whole transformation for a subsystem *i* the fuzzy state assignment 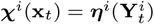.

In practice, we extract all time-lagged data pairs **x**_*t*_, **x**_*t*+*τ*_ that contain all molecular geometry features (e.g., distances, contacts, torsions) of our simulation data and pass them through the architecture presented in Fig. 2. The data is fed through an attention mechanism (represented by the matrix **G**) that yields subsystem specific vectors 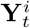, each of which attends to features relevant for subsystem *i*. These vectors then serve as inputs to *N* parallel feature transformations ***η***^*i*^ (parallel VAMPnets) which transform those into output features ***χ***^1^,… ***χ***^*N*^ (with 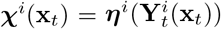) that represent slow collective coordinates or directly fuzzy assignments to metastable Markov states of each molecular subsystem. Equipped with the state assignments, we can compute correlation matrices (Eq. 8) and derive a Koopman model matrix from those (Eq. 9). As in VAMPnets, the feature transformations ***η***^1^,…***η***^*N*^ are represented by deep neural networks. In the present study we use multilayer perceptrons with a SoftMax output layer representing fuzzy state assignments. However, other architectures could be chosen, e.g. graph convolution networks when parameter sharing is desired [67, 68], and a linear output layer could be chosen if the aim is to represent slow collective variable rather than discrete states [57, 58]. The parameters of the feature transformations ***η*** and the attention matrix are learned end-to-end via backpropagation.

In more detail, given N individual subsystem models, the global system state can be given by the Kronecker product of all subsystem states:

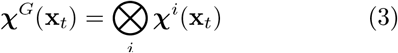

and by computing the global correlation matrices 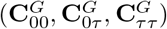 from Eqs. (8) using ***χ***^*G*^. We note that this step does not require that we have independent Markovian models, but it is simply a formalism to express global states in terms of a combination of local states.

Furthermore, we construct a candidate for the global Koopman model from the subsystem models by combining the individual singular values and vectors with a Kronecker product [64]:

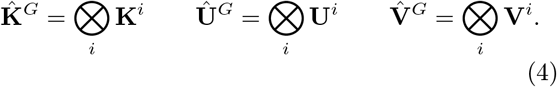

The matrices **Û**^*G*^ and 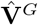 map the global state assignments onto the constructed singular functions and are computed from the local matrices as defined in Eqs. (10–11). The diagonal matrix 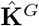 encodes the singular values and is computed from the subsystem singular value matrices via Eq. (9).

In order to evaluate the performance of the constructed model to predict the dynamics in the global state space, the VAMP-E validation [55] score can be exploited,

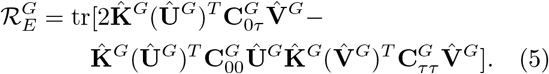

The VAMP-E score measures the difference between the estimated Koopman model and the true dynamics. Here, it is evaluated for the global state assignments 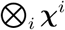 (as encoded in 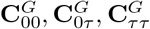) mapped on the constructed singular functions (as encoded in **Û**^G^, 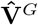). If the subsystems are independent the constructed singular functions are optimal and the singular values of the global system are indeed the product of singular values of the subsystems (as formalized in Conditions for independent systems, also see SI Appendix, Independent Koopman operators). In this case, the global VAMP-E score Eq. (5) has a product form

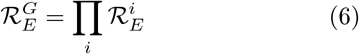

that poses a necessary condition for subsystem independence.

To finally train the model, we develop a loss function that (i) maximizes the global VAMP-E score, assuming that they describe independent dynamics (Eqs. 3–5), and (ii) minimizes a term that penalizes statistical dependence between these subsystems (Eqs. 6) scaled by a weighting factor *ξ*.

We evaluate the scores only pairwise, to escape the growth of the global state space, and sum over all possible pairs *i, j*:

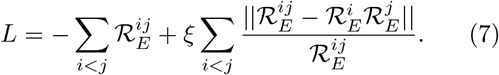

Here, 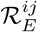 measures the quality of the constructed Koopman model of subsystems *i* and *j* and is computed using Eq. (5). The weighting factor *ξ* is a hyperparameter that should be chosen large enough to find decoupled systems and small enough to not interfer with the subsystem dynamics. Even though the choice of an appropriate *ξ* depends on the nature of the dynamics and the coupling, it is directly related to the training procedure as it, briefly, balances focus of the optimizer between kinetics and decoupling. Further conditions (Eq. 17), which evaluate the independence of the singular functions and values, can be used as post training validation metrics for adjusting *ξ* and for testing to which degree dynamically independent subsystems were found.

## Results

The iVAMPnet architecture, which is implemented using *PyTorch* [69], is depicted in Fig. 2. Generally, various neural network architectures are possible; we here choose fully connected feed forward neural networks with up to 5 hidden layers with 100 nodes each. The scripts to reproduce the results including the details for the training routine, choice of hyperparameters, and network architecture can be found in our GitHub repository. We note that an implementation of VAMPnets is available in the current version of DeepTime [70].

### Benchmark model with two independent subsystems

We first demonstrate that iVAMPnets are capable of decomposing a dynamical system into its independent Markovian subsystems based on observed trajectory data using an exactly decomposable benchmark model (Fig. 3).

**Figure 3:**
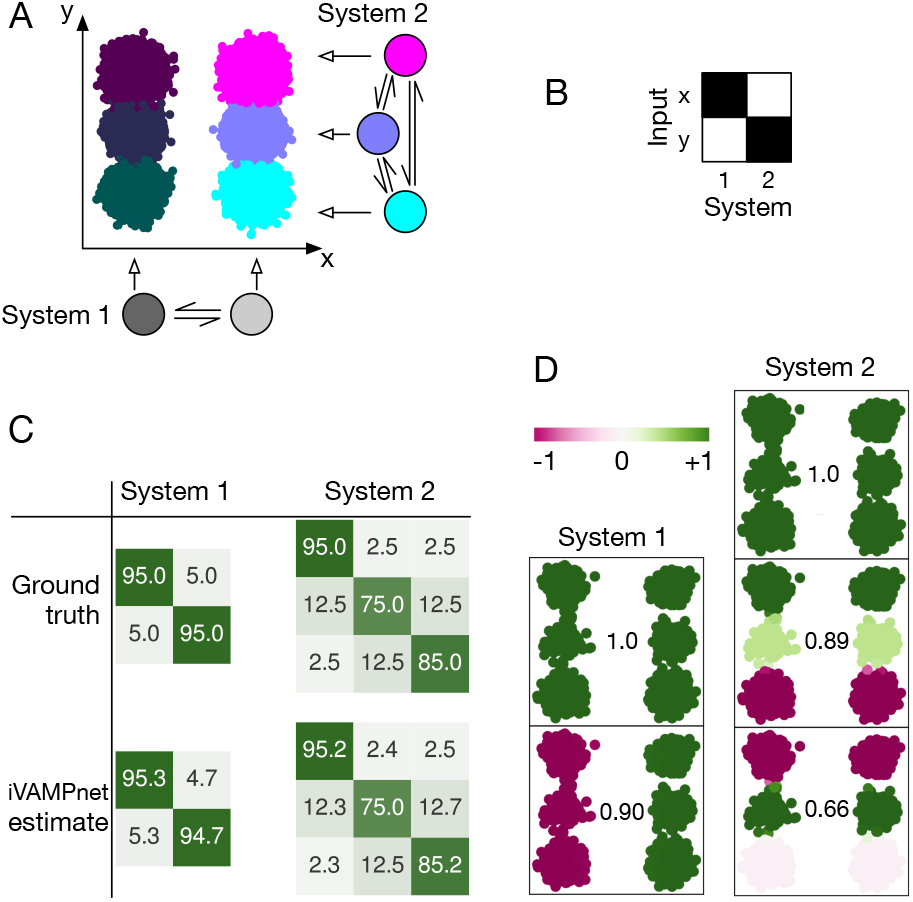
Hidden Markov state model as a benchmark example for independent subsystems: **a)** 2 subsystems with 2 and 3 states emit independently to an *x* and *y* axis, respectively. The corresponding 2D space embeds all 6 global states. **b)** The learned mask shows that each subsystem focuses on one input dimension. **c)** The estimated subsystem transition matrices are compared with the ground truth. **d)** Subsystem eigenfunctions and corresponding eigenvalues as found by iVAMPnet. Independent processes are recovered from the 2D data.

Akin to the protein illustrated in Fig. 1, we define a system that consists of two independent subsystems with two and three states, respectively. It is modeled by two transition matrices with the corresponding number of states. We sample a discrete trajectory with each matrix (100k steps) [70]. The global state is defined as a combination of these discrete states. The discrete subsystem states are now interpreted as the hidden states of hidden Markov models [71] that emit to separate, subsystem-specific dimensions of a 2D space. The output of each subsystem is modeled with Gaussian noise 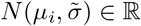 that is specific to the state that the system is in, specified by the mean *μ_i_*, and a constant 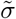. The two state subsystem therefore describes a jump process between Gaussian basins along the *x*-axis and the three state along the *y*-axis, respectively (Fig. 3a). These variables compare to collective variables of the green (*x*) and blue (*y*) system depicted in Fig. 1. Please note that while in this benchmark system the relevant slow collective variables are known, iVAMPnets are generally capable of finding them (cf. 10D hypercube benchmark model and Synaptotagmin-C2A).

Since the generative benchmark model consists of perfectly independent subsystems and the pair already describes the global system, our method can simply be optimized for the global VAMP-E score (Eq. 5) without the need for any further constraints. We train a model with a two and three state subsystem at a lag time of *τ* = 1 step.

Once trained, the iVAMPnet yields a model of the dynamics in each of the identified subsystems. As expected, we find that the estimated transition matrices for both subsystems closely agree with the ground truth (Fig. 3c). To additionally assess the slow subsystem dynamics in more detail, we borrow concepts from MSM analysis and conduct an eigenvalue decomposition of the iVAMPnet models (cf. VAMPnets). The analysis of the eigenfunctions demonstrates that, by construction, the system exhibits one independent process along the *x*-axis (λ_1_ = 0.90) and two along the y-axis (λ_2_ = 0.89 and λ_4_ = 0.66) (Fig. 3d). In contrast, we note that in the picture of *global* states, two additional processes would appear as a result of mixing the independent processes (cf. SI Appendix, Fig. S1), which makes the combined dynamical model more challenging to analyze, whereas the iVAMPnet analysis remains straightforward and simple.

Besides the dynamical models, our iVAMPnet yields assignments between input features and subsystems. We find that the method correctly identifies the two state system as the x-axis and the three states as the y-axis feature, respectively (Fig. 3b).

### 10D hypercube benchmark model

In a next step we test the iVAMPnet approach with ten 2-state subsystems, which corresponds to 1024 global states (Fig. 4a,b). As before, the dynamics is generated by ten independent hidden Markov state models with unique timescales. The system is split into five pairs of subsystems, and the two coordinates governing the transition dynamics of each pair are rotated in order to make them more difficult to separate (Fig. 4a). Additionally, we make the learning problem harder by adding ten noise dimensions such that the global system lives on a 10-dimensional hypercube embedded in a 20 dimensional space.

**Figure 4:**
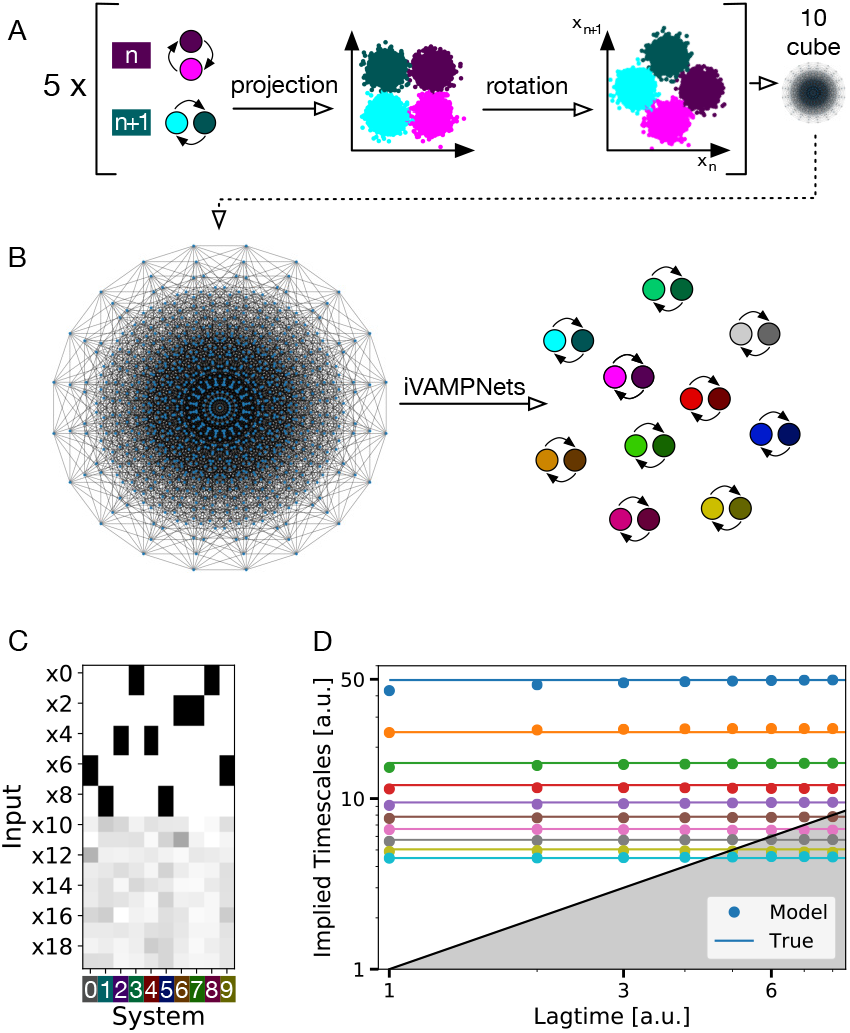
Hidden Markov state model with 1024 global states forming a 10D hypercube embedded in a 20D space. **a)** The hypercube is composed of ten independent 2-state subsystems. A pair of two subsystems always lives in a common rotated 2D-manifold. Therefore, two subsystems need the same input features to be well approximated. **b)** 2D depiction of the hypercube in an orthographic projection [72, 73], where the global system can jump freely between all 1024 vertices, and the ten 2-state models retrieved from it by the iVAMPnet. **c)** Learned mask shows that for each subsystem, the network assigns two highly important input features which are shared with exactly one other subsystem, mirroring the rotated input space. Noise dimensions (x10-x19) are assigned low importance values. **d)** Implied timescales of all ten subsystems learned by our method (dots) approximate the underlying true timescales (lines).

Although the subsystems are perfectly independent, we will estimate an iVAMPnet with the VAMP-E score in a pairwise fashion, thereby avoiding to estimate expensively large correlation matrices in ℝ^1024×1024^. As this is only justified if all systems are independent, we additionally enforce Eq. (6) during training by minimizing Eq. (7) and thereby rule out that any two subsystems approximate the same process.

The iVAMPnet estimation yields subsystem models which, as common in MSM analysis, can be validated by testing whether their implied relaxation timescales are converged in the model lag time *τ*. We find that the implied timescales learned by the iVAMPnet are indeed converged and accurately reproduce the ground truth (Fig. 4d). We note that in addition to the timescales of the individual subsystems that are identified by the iVAMPnet, a global model would also contain all timescales that result from products of eigenvalues, resulting in a total of 1024 timescales. Thus, the iVAMPnet analysis provides a much simpler and more concise model than a global MSM or VAMPnet would.

Furthermore, the subsystem assignment mask indicates that the method correctly assigns high importance weight to two input features for each model (Fig. 4c). Therefore, the method proves its capability of decomposing a noisy, high dimensional global system into its independent sub-processes in a data efficient way.

We have generalized the 10-cube system to a variable number of subsystems (*N*-cube) to conduct a performance benchmark, finding that iVAMPnets outperform VAMPnets for this particular system. We however note that this result may not be generalizable to arbitrary systems as the N-cube features truely independent 2-state subsystems (compare SI Appendix, Performance evaluation for details).

### Synaptotagmin-C2A

Finally, we test iVAMPnets on an all-atom protein system. In comparison to our benchmark examples, we expect the underlying global dynamics to be only approximately decomposable into independent subsystems. Our test data consists of 184 μs aggregate MD data of each 2 μs length (92 × 2 μs) of the C2A domain of synaptotagmin (SI Appendix, MD setups) that was described previously [74]; synaptotagmin plays a crucial role in the regulation of neurotransmitter release [75]. It was shown to consist of approximately uncoupled subsystems containing the calcium binding region (CBR) and the C78 loop, respectively [64].

First, we attempted to model the protein with a global model, i.e., with a single (regular) VAMPnet. Indeed, this approach failed because there were not enough simulation statistics to estimate a reversibly connected transition model between all global metastable states, resulting in diverging implied timescales (SI Appendix, Global model of synaptotagmin and Fig. S2). This is exactly the scenario where iVAMPnets should provide an advantage, by only relying on locally rather than globally converged transition statistics.

Next, we train an iVAMPnet to seek two subsystems of twelve and six states, respectively, each at a lag time of *τ* = 10 ns where we enforce constraint Eq. (6) to find uncoupled subsystems.

The trained iVAMPnet identifies one subsystem comprising all three CBR loops (CBR-1, CBR-2, CBR-3; Fig. 5a). The second subsystem consists not only of the aforementioned C78 loop but also of the loop connecting beta sheets 3 and 4 [76] (termed C34 henceforth). When mapping the residue positions on the protein structure it becomes obvious that the two subsystems are physically well separated (Fig. 5a), supporting the conclusion that both regions are only weakly coupled [64].

**Figure 5:**
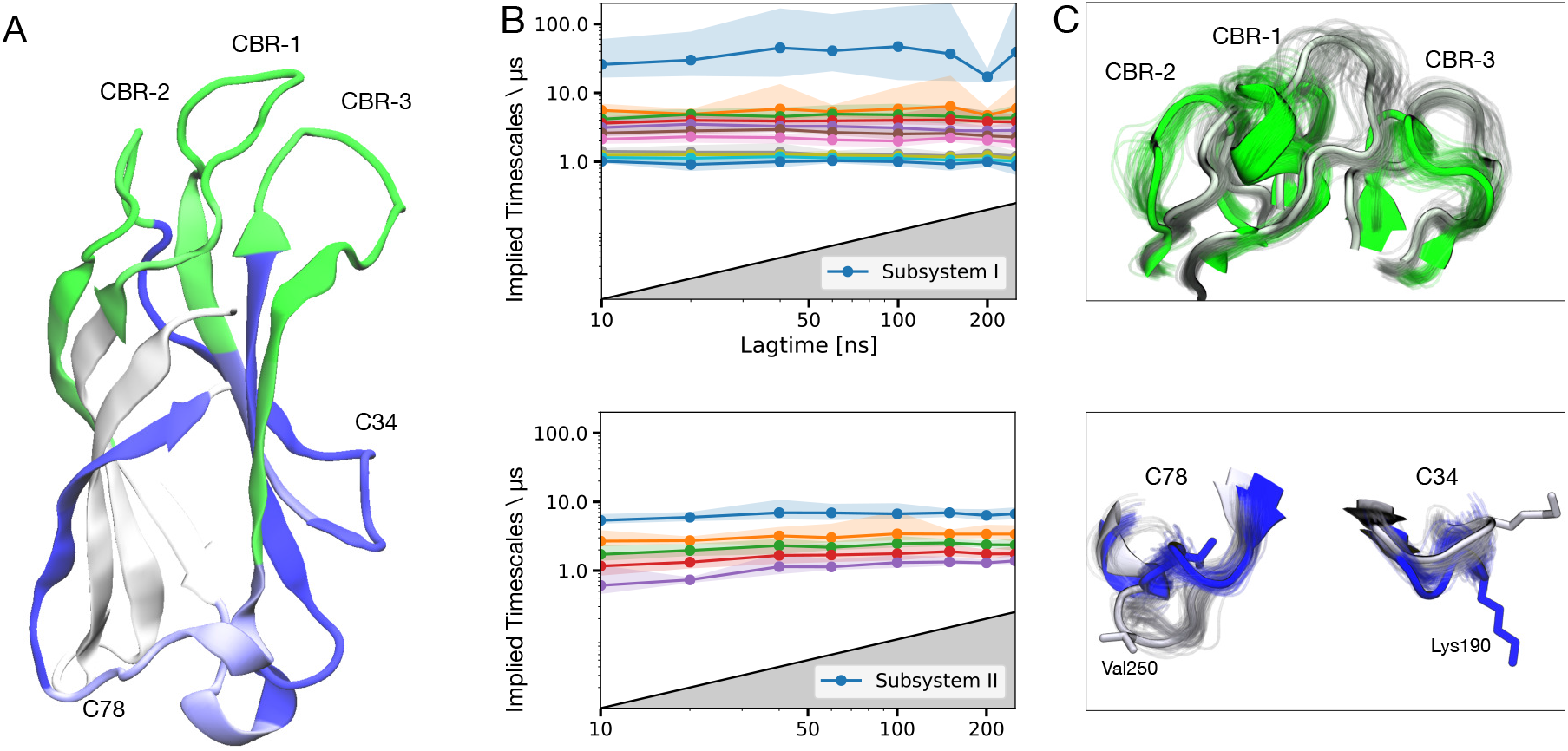
iVAMPnet of synaptotagmin-C2A with two subsystems and twelve and six states, respectively. **a)** Importance values of the trainable mask depicted as color-coded protein secondary structure, indicating assignment to subsystem I (II) in green (blue). **b)** Implied timescales of the two subsystems with a 90% percentile over 20 runs. **c)** Superposed representative structures of both extrema of the slowest resolved eigenfunctions of each subsystem (residues not assigned a high importance value or not showing significant movement are omitted for clarity). The slowest process of subsystem 1 changes between green and gray structures showing an orchestrated movement of the full Calcium Binding Region (CBR1, CBR2, and CBR3). The slowest process of the second subsystem occurs between the blue and gray structures and describes a combined movement of C78 and C34.

The implied timescales of both systems are approximately constant in the model lag time *τ*. Most timescales are in the range of 1 – 10 *μ*s, with the exception of one much slower process with a 100 μs relaxation time found in the first subsystem (Fig. 5b), which has not been found previously. Analysis of the structural changes governing this process reveals that it involves an orchestrated transition of all CBR loops (Fig. 5c). Such a process could however not be resolved by the previous study [74] where the CBR was modeled as individual loops. The process of the second system involves a simultaneous movement of the C78 and C34 loops (Fig. 5c).

iVAMPnets find metastable structures in the local features that are comparable to the ones described in our previous work [74]. Specifically, α-helices in two distinct positions and a state burying a methionine residue (Met173) can be found in the CBR1. In the adjacent CBR2 site, both tightly bound and loose configurations are identified, and the C78 site features all three previously described valine residue conformations (Va1250, Va1255). In addition to the features modeled in our preceding study [74], iVAMP-nets identify dynamics in a lysine rich cluster (Lys189-192) that was previously reported as important for membrane interaction [77]. Please compare SI Appendix, iVAMPnet model of Syt-C2A for a detailed view on the metastable states and exchange kinetics. In contrast to our previous work, the kinetic models in the local subsystems are more complex and incorporate a larger number of dynamic processes, providing a more comprehensive picture without the need to define a partitioning manually. In fact, conducting domain-decomposition and local kinetic modeling simultaneously has enabled the identification of very subtle dynamical features as long as they contribute significantly to the local VAMP-scores.

Although estimating a global VAMPnet model for synaptotagmin was not feasible given the sparse data sample, iVAMPnets use the same data efficiently and estimate a statistically valid dynamical model. This result is especially striking because the iVAMPnet approach also simplifies the subsequent task of interpreting models by separating dynamically independent protein domains.

### Counterexample: folding of the villin miniprotein

Finally, we conducted an experiment on a villin protein folding trajectory of 125 μs length [78] as a negative example (SI Appendix, MD setups). Small proteins such as villin are typically cooperative, i.e., the slowest processes related to folding involve all residues (SI Appendix, Villin: A non-decomposable system). Thus, these processes cannot be resolved when decomposing the system into several subsys-tems. Indeed, we find that a splitting into two subsystems with two states each results in timescales that are not converged, and whose relaxation processes approximate a partial folding on disjoint areas (cf. SI Appendix, Fig. S6).

### Testing statistical independence of the learned dynamical subsystems

As constraint Eq. (6) was used as a penalty during training (as independence score Eq. (18)), we assess the validity of an estimated subsystem assignment by evaluating the constraints that were not enforced during training (Eq. 16) as post-training independence scores *M_U_,M_V_*, and *M_UV_* (defined in Eq. (17)). Low values for *M_U_* and *M_V_* imply that the constructed left and right singular functions are indeed valid candidates for singular functions in the global state space. A small value for *M_UV_* indicates that the kinetics in the global state space is well predicted by the Kronecker product of subsystem models. We find that the three metrics are well suited to indicate independence of the learned subsystems (Tab. 1). Out of the tested systems only villin cannot be split into independent parts (all scores > 0.1). In comparison, the benchmark models and synaptotagmin can be decomposed into statistically uncoupled subsystems (all scores < 0.01). The slightly increased *M_R_*-value for synaptotagmin suggest that its subsystems might be weakly coupled.

**Table 1:** Post-training independence validation. The scores in columns 1-3 (*M_U_*, *M_V_*, *M_UV_*, cf. Eq. (17)) are computed from independence constraints that were not enforced during the training. The score in the last column (*M_R_*) is used during the training and shown for reference. The three post-training validation scores *M_U_,M_V_*, and *M_UV_* indicate that the final subsystems of both benchmark examples and synaptotagmin are indeed independent, whereas the scores for villin strictly oppose this conjunction. The standard deviations (SD) over 10 different runs are on the order of 10^-5^ for all systems except villin, which has an SD ~ 10^-4^.

| | $M_U$ | $M_V$ | $M_{UV}$ | $M_R$ |
| --- | --- | --- | --- | --- |
| Benchmark 2 | 0.0058 | 0.0059 | 0.0055 | 0.0002 |
| 10-Cube | 0.0039 | 0.0039 | 0.0046 | 0.0005 |
| Synaptotagmin | 0.0042 | 0.0042 | 0.0044 | 0.0018 |
| Villin | 0.1353 | 0.1364 | 0.1493 | 0.0021 |

## Discussion

We have proposed an unsupervised deep learning framework that, using only molecular dynamics simulation data, learns to decompose a complex molecular system into subsystems which behave as approximately independent Markov models. Thereby, iVAMPnet is an end-to-end learning framework that points a way out of the exponentially growing demand for simulation data that is required to sample increasingly large biomolecular complexes.

Specifically, we have developed and demonstrated iVAMPnets for molecular dynamics, but the approach is, in principle, also applicable to different application areas, such as fluid dynamics. The specific implementation, such as the representation of the input vectors **x**_*t*_ and the neural network architecture of the ***χ***-functions, depend on the application and can be adapted as needed.

We now have a hierarchy of increasingly powerful models ranging from MSMs over VAMPnets to iVAMPnets. MSMs always consist of (1) a state space decomposition and (2) a Markovian transition matrix governing the dynamics between these states. VAMPnets provide a deep learning framework for MSMs, and thereby (3) learn the collective coordinates in which the state space discretization (1) is best made. iVAMPnets additionally learn (4) a physical separation of the molecular system into subsystems, each of which has its own slow coordinates, Markov states, and transition matrix.

We have demonstrated that iVAMPNets are a powerful multiscale learning method that succeeds in finding and modeling molecular subsystems when these subsystems indeed evolve statistically independently. Additionally, iVAMPnets are capable of learning from high dimensional MD data. To prove that point, we have demonstrated that the synaptotagmin C2A domain is decomposable into two almost independent Markov state models. Importantly, we have shown that this dynamical decomposition of synaptotagmin C2A succeeds while an attempt to model the system with a global Markov state model fails due to poor sampling. This is a direct demonstration that iVAMPnets are statistically more efficient than VAMPnets, MSMs or other global-state models and may indeed scale to much larger systems.

We note, however, that iVAMPnets do not learn how the subsystems are coupled, and are therefore, in their current form, only applicable to molecular systems that consist of uncoupled or weakly coupled subsystems. Although most biomolecular complexes are known to be cooperative, there are examples that have been modeled very successfully using independent subsystems, such as the Hudgkin-Huxley model of voltage gated channel proteins [79, 80]. For other systems, the degree of coupling is a matter of debate, for example the C2-tandem (C2A and C2B domains) in synaptotagmins [81, 82]. Since isolated domains are known to conduct function by themselves in many cases, we believe that discarding couplings is a first-order modeling assumption that is suitable to identify these domains and their relevant metastable states.

Following up on Ref. [63] and introducing coupling parameters that describe how the learned MSMs are coupled, is subject to ongoing research. Furthermore, the weak-coupling assumption is made for the timescale of the investigated molecular processes and may not be generalizable to arbitrary times. E.g., the degree of coupling between domains found in an MD simulations of a folded protein state may be very different in its unfolded state, which will be eventually encountered for a long enough simulation time.

Besides the usual hyperparameter choices in deep learning approaches, iVAMPnets require the specification of the number of sought subsystems. This choice can be guided by training an iVAMPnet for different numbers of subsystems and then interrogating the independence scores (Eq. 18 and Eqs. 17) to choose a decomposition where statistical independence is optimal. We suggest to start with decomposing the system into two subsystems as a starting point, and to increase this number subsequently. Non-optimal choices may, e.g., reflect in nonconverged implied timescales (possibly an incarnation of the sampling problem that may be mitigated by increasing the number of subsystems) or high independence scores (not possible to split the system because too many or non-optimal number of subsystems were chosen). Furthermore, the choice of the number of subsystems can be guided by the number of structural domains in a protein (complex) or by using the network-based approach presented in Ref. [64]. Furthermore, the number of states in each subsystems needs to balance a) the quality of the singular function approximation (higher for few states) and b) model resolution (higher for more states). Ultimately, different choices may yield converged validation measures, and the number of states may be chosen to yield the desired model resolution in this case.

iVAMPnets can be improved and further developed in multiple ways, e.g. by employing more advanced network architectures, e.g. graph neural networks, where parameters could be shared across subsystems. This might result in higher quality models and a greater robustness against the hyperparameter choice. Very recently, graph neural networks were indeed successfully combined with VAMPnets, showing that the resulting method (GraphVAMPnets) is applicable to MD data and that the estimated models are high quality [83].

In summary, iVAMPnets pave a possible path for modeling the kinetics of large biological systems in a data-efficient and interpretable manner.

## Methods

### VAMPnets

Since an iVAMPnet implements multiple parallel VAMPnets representing the kinetics of separate independent subsystems, we will introduce VAMPnets first [56]. VAMPnets are multilayer perceptrons that represent feature functions ***χ*** (we omit the upper subsystem index *i* for the sake of clearness here). Their last layer is often chosen to be a SoftMax function, i.e., summing over all non-negative outputs yields a 1. Therefore, the output of a VAMPnet can be interpreted as a fuzzy assignment to a metastable state. Taking the linear combination of states with equal weights results in the constant singular function with the singular value 1, which will be reflected by the singular values of the Koopman matrix (Eq. 9 with the normalized correlation matrix). Given the feature functions ***χ***, we can compute the following correlation matrices:

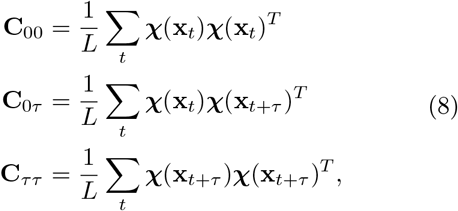

where *L* is the number of collected data pairs in the simulations.

Training VAMPnets or iVAMPnets involves the computation of covariance matrices over minibatches. We therefore need to choose the batchsize to balance large estimator variance obtained for small batches and high memory requirements for large batches. Instead of using the trivial covariance estimator (Eqs. 8) which is asymptotically unbiased [55] but has a high-variance, one can employ a shrinkage estimator [84, 85] which reduces the overall estimator error by trading larger bias for lower variance. For the current study, we assume that our benchmark and MD data has been sufficiently sampled to yield adequate approximations with the estimator given in Eqs. 8.

The approximation of the singular functions and values can be estimated via the singular value decomposition (SVD) of the following matrix 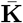:

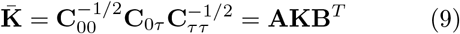

**K** is the diagonal matrix of approximated singular values corresponding to the left and right singular functions:

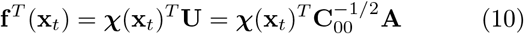

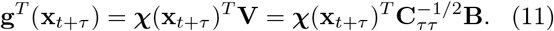

The matrices **U** and **V** construct the left and right singular functions from the individual state assignments. The optimal state assignments can be found by maximizing the VAMP-E score:

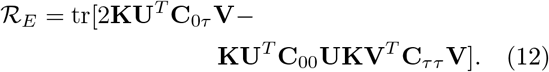

Given trained state assignments ***χ***(**x**_*t*_) and correlation matrices Eq. (8), the Koopman matrix **T** can then be evaluated as:

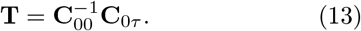

Further, we can estimate the eigenfunction ***φ*** and timescales *t_i_* by its eigendecomposition **T** = **QΛQ**^-1^:

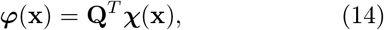

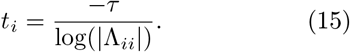

Please note that this operation is only possible if the eigendecomposition is (approximately) real-valued, a condition that is met for the presented application cases.

### Conditions for independent systems

For Markov independent systems, the singular values and functions that are constructed by the Kronecker product match the true global ones,

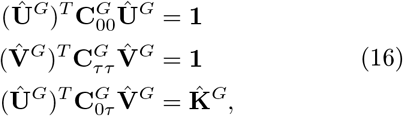

where the first two equations guarantee the orthonormality of the constructed singular functions. The latter verifies that the left and right singular functions correlate as predicted by the Kronecker product of the singular values.

These conditions can be translated to the following scores:

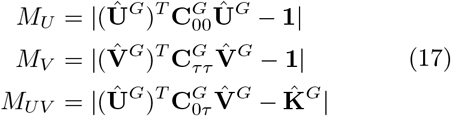

Furthermore, using the identities Eq. (16) and the definition of the VAMP-E score Eq. (12) yields

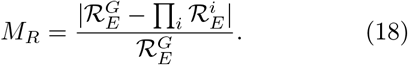

The norms denote simple means. The last score, *M_R_*, is enforced during training in a pairwise fashion (cf. Eq. 7).

### Network architecture

Given a global system, which we want to decompose into *N* subsystems, and a time series of input features 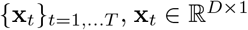, we pass the features through a mask **G** ∈ ℝ^*D*×*N*^, which weights each input differently for each subsystem, before the result are transformed individually by the *N* independent state assignment functions ***η***^*i*^. It should be mentioned that the mask is merely introduced for interpretability reasons and is not essential to find independent subsystems. If the mask was omitted, the extraction of the relevant features would simply be transferred to the downstream neural networks, remaining hidden to the practitioner.

The weighted input is assessed by an element wise multiplication 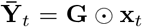. In order to prevent the neural networks to reverse the weighting of the mask in its consecutive layers, we draw for each input feature i and subsystem *j* an independent, normally distributed random variable 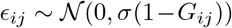. This noise is added to the weighted features:

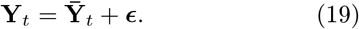

Thereby, the attention weight linearly interpolates between input feature and Gaussian noise, i.e., if the attention weight *G_ij_* = 1, *Y_ij_* carries exclusively the input feature *x_i_*, if *G_ij_* = 0, *Y_ij_* is simple Gaussian noise. By tuning the noise scaling *σ*, a harder assignment by **G** can be enforced. This hyperparameter should be optimized by adjusting it so that the resulting mask yields clear subsystem assignments without being binary. Subsequently, the transformed feature vector is split for each individual subsystem 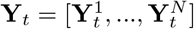 and passed through the subsystem specific neural network ***η***^*i*^ resulting in feature transformations 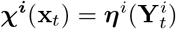. These features are then used to estimate the Koopman models.

### Constructing the mask

To train an interpretable mask, we use the following three premises:

1. A single subsystem should not focus on all input features.
2. Different subsystems compete for high weights for the same feature.
3. All weights should be in the range [0,1] and the matrix should be sparse.

Therefore, the mask is constructed by trainable weights **g** ∈ ℝ^*D*×*N*^ which are first processed by a softmax function which normalizes along the input feature axis **g**_1_ = softmax(**g**, dim=0). Thereby, if a subsystem focuses on one part of the features, a lower weight for the other parts is expected following the ñrst premise.

In a next step, weights which are lower than a threshold *θ* are clipped to zero **g**_2_ = relu(**g**_1_ – *θ*) to guarantee sparsity. The threshold θ is a hyperparameter that can be optimized by starting with comparably small values (i.e., very little cutoff) and subsequently increasing it without further training – a reasonable cutoff does not alter the results in this case, as the downstream neural networks still obtain all relevant information.

Since input features could be negligible for all subsystems, a dummy system is added which has a constant value **c** ∈ ℝ^*D*×1^ for all features **g**_3_ = [**g**_2_, **c**]. Consequently, the weights of all subsystems and the dummy system are normed for each feature **g**_4_ = **g**_3_/sum(**g**_3_, dim = 1), which together with the clipping fulfills the premises two and three.

Finally, the mask is given by truncating the dummy system 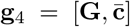. Beware that only **g**_4_ is normalized along the system axis.

### Application to protein dynamics

Since for proteins the final model is often expected to be invariant with respect to rotations and translations, internal coordinates are employed as input features. For Markov state modeling, the minimal heavy atom distance *d_ij_* between residues *i, j* has been proven to be a good descriptor [56, 86]. However, for interpretability, mask weights for each residue are preferable. Therefore, the mask is of size **G** ∈ ℝ^*R*×*N*^ with the number of residues *R*. The input features are then scaled as *x_ij_ G_i_G_j_* exp(−*d_ij_*).

Furthermore, a smoothing routine is implemented such that neighboring residues along the chain have similar importance weights. *W* windows of size *B* are placed along the chain with step size *s*. Each window has a trainable weight **g** ∈ ℝ^*W*×*N*^. Consequently, the softmax function is taken along the window axis 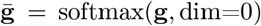. However, before applying the clipping as before the weight for each residue **g**_1_∈ ℝ^*R×N*^ is calculated as the product of all window weights the residue is part of (Fig. 6).

**Figure 6:**
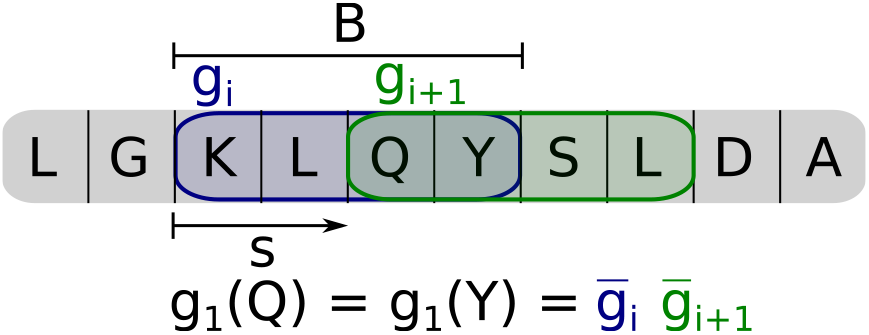
Attention scheme for amino acid chain. Windows of size *B* are placed along the chain with a step size of *s* resulting into *W* many windows. A trainable weight **g** ∈ ℝ^*W*×*N*^ is assigned for a window in each subsystem which are made positive and normalized along the window axis through a softmax 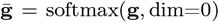. Here a window size of *B* = 4 and a step size of *s* = 2 is chosen. As a consequence the weight of the amino acid glutamine (Q) is given as the product of the two windows it is part of 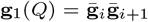. The choice of the step size determines how many neighboring amino acids have the exact same weight within a subsystem, which applies here for the tyrosine (Y). Together with the window size it is regulated how many residues share parts of their weights. Hence, the serine (S) shares the weight 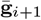 with the previous two amino acids 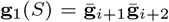, which has a smoothing effect on the attention mechanism along the chain.

## Data availability

The benchmark data can be generated from the Jupyter notebooks published in our GitHub repository. The molecular dynamics data set of synaptotagmin C2A is available at https://doi.org/10.5281/zenodo.6908073. Restrictions apply to the availability of the villin data set, which were used under license for this study. Data are available from the authors upon request. The villin headpiece folding data is courtesy of D.E. SHAW research [78].

## Code availability

The code that implements the presented models and reproduces the presented results can be found in our GitHub repository.

## Ackowledgements

We acknowledge financial support from Deutsche Forschungsgemeinschaft DFG (SFB/TRR 186, Project A12; SFB 1114, Projects C03 and A04; SFB 1078, Project C7, and RTG 2433), the European Commission (ERC CoG 772230 “ScaleCell”), the Berlin Mathematics center MATH+ (AA1-6 and AA1-10), the BMBF (Research center BIFOLD), the National Science Foundation (CHE-1738990, CHE-1900374, and PHY-2019745), the Welch Foundation (C-1570), and the Einstein Foundation Berlin. We further thank Manuel Dibak and Moritz Hoffmann (FU Berlin) for fruitful discussions.

## Notes

**Conflicts of interest:** The authors declare no conflicts of interests.

https://github.com/markovmodel/ivampnets

### Competing Interest Statement

The authors have declared no competing interest.

### Summary of Updates

Revision clarifying open questions and contains in-detail analysis of Syt-C2a iVAMPnet model.

