## Supplemental material for "Deep learning to decompose macromolecules into independent Markovian domains"

#### Independent Koopman operators

The true Koopman operator $\mathcal{K}_\tau$ can be written in two ways. First, if the operator is a Hilbert-Schmidt operator, the following singular value decomposition (SVD) exists:

$$\mathcal{K}_\tau g(\mathbf{x}) = \sum_{i=1}^{\infty} \sigma_i \langle g, \phi_i \rangle_{\rho_1} \psi_i(\mathbf{x}). \tag{1}$$

A low-rank approximation to the Koopman operator is obtained by truncating the sum after $k \ll \infty$ terms.

Second, the Koopman operator can be expressed via the transition density $p_\tau(\mathbf{y}|\mathbf{x})$ that describes the transitions from configuration $\mathbf{x}$ to $\mathbf{y}$ within a time window $\tau$:

$$\mathcal{K}_\tau g(\mathbf{x}) = \int p_\tau(\mathbf{y}|\mathbf{x}) g(\mathbf{y}) \mathrm{d}\mathbf{y}. \tag{2}$$

Given two independent systems with configurations $\mathbf{x}^1, \mathbf{x}^2$ and $\mathbf{y}^1, \mathbf{y}^2$ and their transition densities $p_\tau^1(\mathbf{y}^1|\mathbf{x}^1)$, $p_\tau^2(\mathbf{y}^2|\mathbf{x}^2)$, respectively, the global transition density is then:

$$p_\tau^G(\mathbf{y}^1, \mathbf{y}^2|\mathbf{x}^1, \mathbf{x}^2) = p_\tau^1(\mathbf{y}^1|\mathbf{x}^1) \cdot p_\tau^2(\mathbf{y}^2|\mathbf{x}^2). \tag{3}$$

As a consequence, if we study observables which can be expressed as the product of the subsystem specific observables (hence the Kronecker product of the feature functions $\boldsymbol{\chi}$) $g^G(\mathbf{x}^1, \mathbf{x}^2) = g^1(\mathbf{x}^1) g^2(\mathbf{x}^2)$, the global Koopman operator decomposes into the product of subsystem operators:

$$\begin{aligned} \mathcal{K}_\tau^G g^G(\mathbf{x}^1, \mathbf{x}^2) &= \iint p_\tau^G(\mathbf{y}^1, \mathbf{y}^2|\mathbf{x}^1, \mathbf{x}^2) g^G(\mathbf{y}^1, \mathbf{y}^2) \mathrm{d}\mathbf{y}^1 \mathrm{d}\mathbf{y}^2 \\ &= \int p_\tau^1(\mathbf{y}^1|\mathbf{x}^1) g^1(\mathbf{y}^1) \mathrm{d}\mathbf{y}^1 \int p_\tau^2(\mathbf{y}^2|\mathbf{x}^2) g^2(\mathbf{y}^2) \mathrm{d}\mathbf{y}^2 \\ &= \mathcal{K}_\tau^1 g^1(\mathbf{x}^1) \mathcal{K}_\tau^2 g^2(\mathbf{x}^2). \end{aligned} \tag{4}$$

The low rank approximation of the decomposed Koopman operator (defined in Eq. 1) for independent systems can now be written as follows, taking into account only $k^1$ and $k^2$ singular functions:

$$\begin{aligned} \hat{\mathcal{K}}_\tau^G g^G(\mathbf{x}^1, \mathbf{x}^2) &= \hat{\mathcal{K}}_\tau^1 g^1(\mathbf{x}^1) \hat{\mathcal{K}}_\tau^2 g^2(\mathbf{x}^2) \\ &= \sum_{i=1}^{k^1} \sigma_i^1 \langle g^1, \phi_i^1 \rangle_{\rho_1^1} \psi_i^1(\mathbf{x}^1) \sum_{j=1}^{k^2} \sigma_j^2 \langle g^2, \phi_j^2 \rangle_{\rho_1^2} \psi_j^2(\mathbf{x}^2) \\ &= \sum_{i=1}^{k^1} \sum_{j=1}^{k^2} \sigma_i^1 \sigma_j^2 \langle g^1 g^2, \phi_i^1 \phi_j^2 \rangle_{\rho_1^G} \psi_i^1(\mathbf{x}^1) \psi_j^2(\mathbf{x}^2) \\ &= \sum_{l=1}^{k^1 k^2} \sigma_l^G \langle g^G, \phi_l^G \rangle_{\rho_1^G} \psi_l^G, \end{aligned} \tag{5}$$

Therefore in case of independent systems, the optimal singular functions and values of the *global* system are given by the Kronecker product of the *subsystem* singular functions and values, $\sigma_l^G = \sigma_i^1 \sigma_j^2$, $\psi_l^G = \psi_i^1 \psi_j^2$, and $\phi_l^G = \phi_i^1 \phi_j^2$. This procedure can be applied to arbitrarily many independent subsystems.

### Global model of 3x2 benchmark system

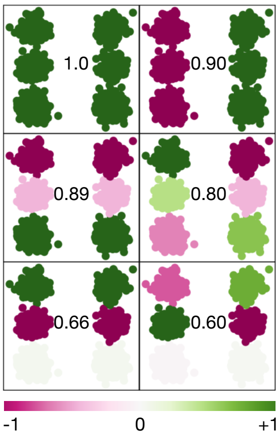


**Figure S1:** Hidden Markov state model as a benchmark example for independent subsystems: The 6 global eigenfunctions supplied with their eigenvalues revealing the 4 independent processes and the 2 resulting mixed product processes. Eigenvalues of the latter are computed from the product of independent process eigenvalues: $\lambda_4 = \lambda_2 \cdot \lambda_3 = 0.80$ and $\lambda_6 = \lambda_2 \cdot \lambda_5 = 0.60$

### Global model of Syt-C2A

To compare iVAMPnets to existing methods, we estimate a classical (global) VAMPnet model with 8 output nodes and no attention mechanism, but otherwise the same hyperparameters that were used for the iVAMPnet estimation (lag time, batch size, architecture, and training routine).

The training score converges to the theoretical maximum of 8. However, when projecting on the eigenfunctions, it becomes apparent that some of them do not describe any real transition event, i.e., stay constant during each single trajectory. Rather, these eigenfunctions model disconnected configurations that belong to different trajectories, which is an artifact of sparse sampling seeded from multiple distinct configurations. It manifests in the implied timescales (Fig. 2) that become infinite for these processes, i.e., the eigenvalues are $\approx 1$, resulting in numerically unstable implied timescales calculations.

These results imply that in order to model the global Syt-C2A system with classical VAMPnets, more simulation data has to be collected to connect these structures, i.e., the amount of data is not sufficient to build a global model. The iVAMPnet model does not suffer from these shortcomings because products of the mentioned disconnected processes will not be observable in the data, which would result in a lower score. Therefore, the VAMP-E score will favor the processes which are truly observed.

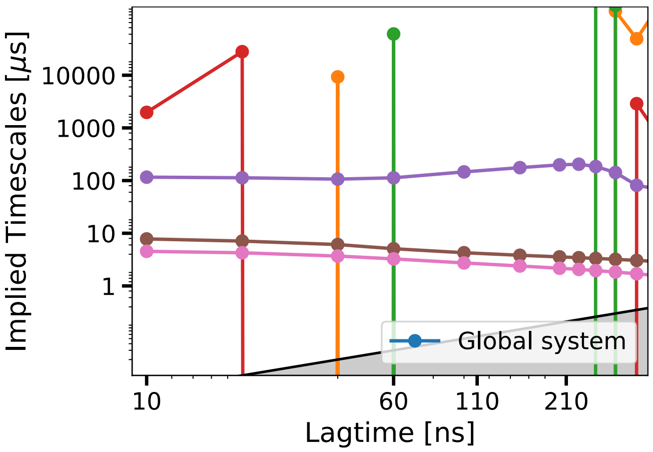


**Figure S2:** Failed global implied timescales test with a classical VAMPnet of synaptotagmin with 8 output nodes. Since the model resolves processes which are not connected, the eigenvalues are $\approx 1$, making the implied timescales estimation numerical unstable. It indicates that the amount of data is insufficient to build a global model. The model has no attention mechanism, but utilizes otherwise the same hyper-parameter than the ones used in iVAMPNets.

### iVAMPnet model of Syt-C2A

The iVAMPnet model of synaptotagmin presented in this paper identifies two distinct subsystems that roughly correspond to the calcium binding region (CBR) and the opposite site of the protein, in particular, two loops that we call C34 and C78. In the following, we discuss the features of our iVAMPnet models that describe these subsystems individually. To that end, we make use of the system description provided in our earlier publication [1]. Furthermore, we approximate transition probabilities and stationary probabilities by computing the transition operator $C_{00}^{-1}C_{0\tau}$ using the iVAMPnet basis functions in each of the subsystems. We note that this approximate transition model is given only for orientation and comparison with the previous model, since the approximated transition operator is generally not a row-stochastic matrix (also compare [2]).

In general, we find that the structural features resolved by our previous study are also resolved by iVAMPnets. In the CBR (cf. Table 1), we find CBR1 $\alpha$-helices at two locations and a state burying Met173, as well as a structural rearrangement in the CBR2 that may be described as attachment / detachment of that loop to the protein body. Furthermore, iVAMPnets identify metastable dynamics in the CBR3 loop that was not previously described. In comparison to our previous model, the iVAMPnet model for the CBR describes kinetics involving concerted motions of all three CBR loops. We show structural renders of the metastable states in Fig. 3.

The second subsystem identifyed by iVAMPnets contains the C34 and C78 loops (Fig. 4): The C78 loop shows structural features as described in [1], i.e., loop rearrangements that are governed by different conformations of two valine residues (Val250, Val255) (Tab. 2). However, iVAMPnets identify another related site that we call C34, a region that is rich in lysine and has been previously described as important for membrane interactions [3]. Its metastable states are described by different conformations of three lysines (Lys 189-191).

Finally, we compare the probabilities of our previous model [1] with the approximate stationary probabilities obtained here (Fig. 5) and find that both models agree qualitatively. Differences between stationary probabilities may be due to the previous model's non-optimal subsystem decomposition and differences in metastable state assignments, in particular regarding the CBR1.

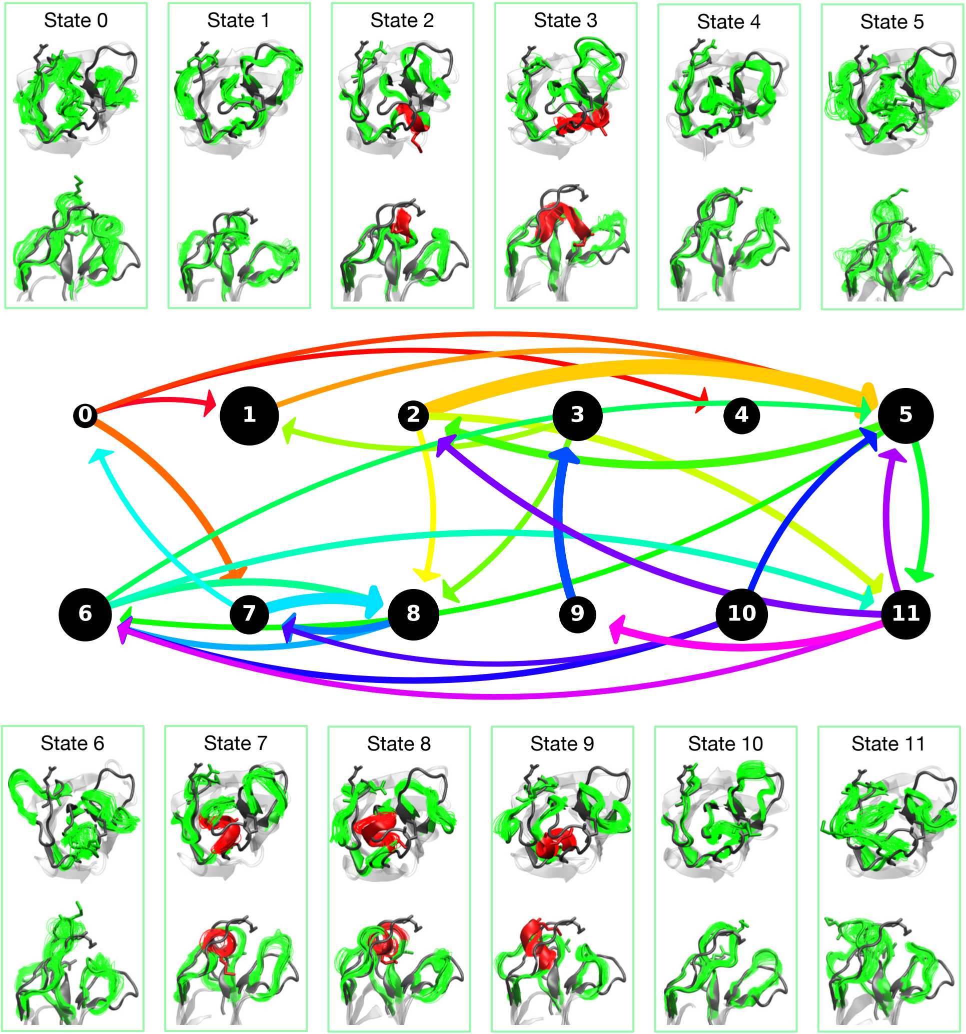


**Figure S3:** Metastable states and transition model of the first subsystem of synaptotagmin that is located in the CBR. For each state, structural renders from two perspectives are given. Helical conformations are highlighted in red. The transition model is an approximation; we show only the most important pathways ($T_{ij} \geq 0.0008$) and depict transition probabilities by arrow thickness. Additionally, approximated stationary probabilities are shown as circle diameters for each state. Arrows are colorized to make them distinguishable.

| **iVAMPnet state** | **legacy CBR1** | **legacy CBR2** |
|---|---|---|
| 0 | disordered (D) | in (A) |
| 1 | Met-in (B) | out (B) |
| 2 | disordered (D) | in (A) |
| 3 | disordered (D) | in (A) |
| 4 | disordered (D) | in (A) |
| 5 | disordered (D) | out (B) |
| 6 | disordered (D) | n/a |
| 7 | top-helix (A) | in (A) |
| 8 | top-helix (A) | out (B) |
| 9 | site-helix (C) | out (B) |
| 10 | disordered (D) | in (A) |
| 11 | site-helix (C)* | out (B) |

**Table S1:** iVAMPnet states of the first subsystem (CBR, shown in Fig. 3) and their correspondence to our previous model ("legacy") [1]. *State 11, which is structurally similar to state 9, was assigned to legacy state C to incorporate uncertainties of the metastable state assignment in our previous model.

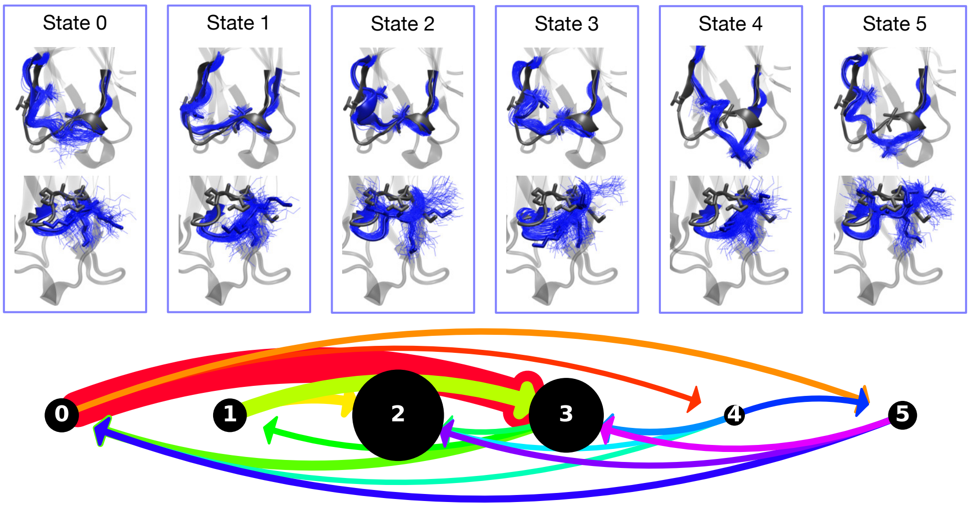


**Figure S4:** Metastable states and transition model of the second subsystem of synaptotagmin, which corresponds to the site opposite of the CBR (loops C78 and C34). Each state is depicted by two structural renders, showing details of C78 (top) and C34 (bottom), respectively. The transition model is explained in the caption of Fig. 3.

| **iVAMPnet state** | **legacy C78** |
|---|---|
| 0 | C |
| 1 | A |
| 2 | C |
| 3 | C |
| 4 | n/a |
| 5 | B |

**Table S2:** iVAMPnet states of the second subsystem (C34 and C78, shown in Fig. 4) and their correspondence to our previous model ("legacy") [1].

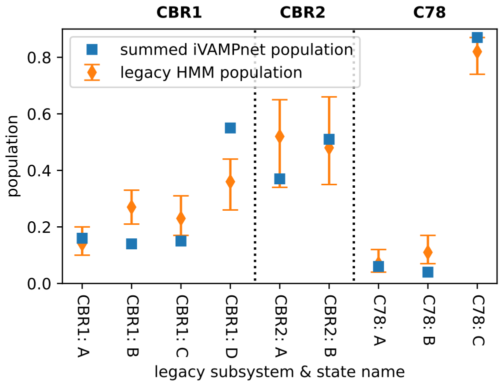


**Figure S5:** Comparison of approximated stationary probabilities of iVAMPnet states with estimates of hidden Markov models (HMMs) given in Ref. [1]. iVAMPnet populations are obtained by summing probabilities over all states that are in a certain legacy state (cf. Tabs. 1,2), the shaded area represents HMM model uncertainty (5-95% percentile of Bayesian HMM samples that were obtained using a mixed prior [4, 5], cf. Ref. [1] for details).

#### Counter-example villin: A non-decomposable system

We demonstrate the behavior of iVAMPnets when confronted with a system that cannot be decomposed into subsystems without missing the slowest global processes. Therefore, we employ the method on the villin dataset [6] where several folding and unfolding events are encoded. If studied by a classical VAMPnet, the folding is recovered as the second slowest process. The slowest process describes a transition between a mis-folded and the folded state [7]. However, the majority of residues are involved in both of these processes, making it hard to decouple these processes into independent subsystems.

For the analysis, the same hyperparameters are used as for synaptotagmin, but we choose only two states per subsystem. The resolved processes resemble a localized folding of either the left or right helix of the folded structure in each subsystem (Fig. 6). However, the implied timescales do not converge, expressing non-Markovian behavior. The results can be interpreted as representing a compromise between learning nearly independent processes and approximating slow processes.

Since the independence is strongly enforced, the processes are badly approximated, resulting in unconverged implied timescales. In order to interpret this model, we have furthermore correlated the iVAMPnet eigenfunctions with the ones of a standard global VAMPNet (Fig. 7). We find that the process found by subsystem 1 has the highest Pearson correlation ($r = -0.79$) with the second global process, which corresponds to peptide folding. However, the second subsystem cannot be clearly assigned to a global process. These results are not surprising since the poor implied timescales convergence and the post-training validation scores (Tab. 1) indicate that the independence approximation does not hold in this example. The system expresses dynamics on the global level that are not or are only poorly approximated by the described iVAMPnets.

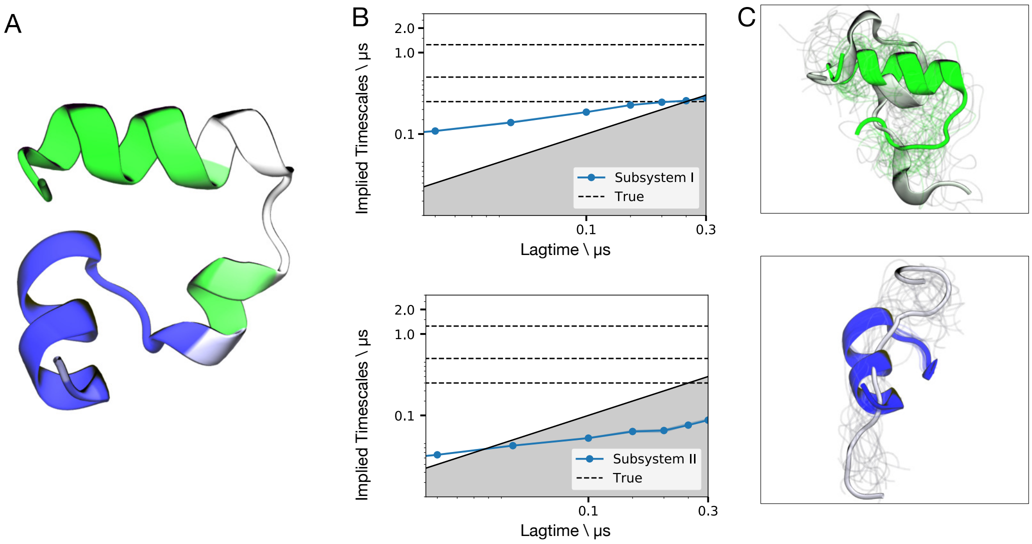


**Figure S6:** Counter example to iVAMPNets using villin, trained with independence constraint. a) Subsystem assignment, i.e., masked importance values, are shown as color code on the folded structure. b) Implied timescales of the 2 subsystems, the black dotted lines are reference timescales of a global model trained with a standard VAMPnet. c) 20 representative structures of both extrema of the slowest resolved eigenfunctions for both subsystems. The processes tend to approximate formation of the N- and C-terminal helices, respectively.

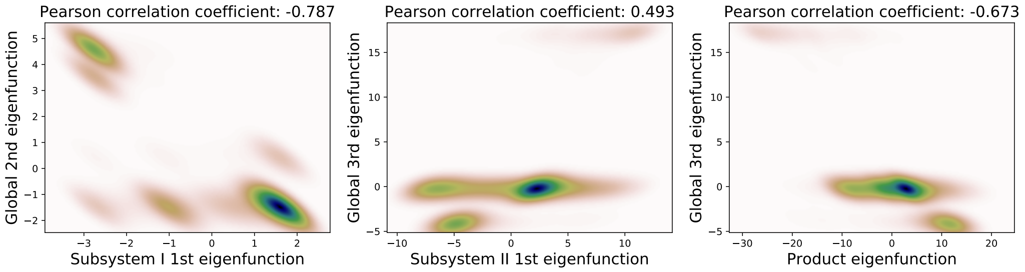


**Figure S7:** Interpreting the processes found by the iVAMPnets for villin by correlating them with the global processes found with a standard VAMPnet. Shown are the processes with the highest correlation plotted against each other. The first subsystem correlates with the second global process, which relates to peptide folding. The second subsystem cannot clearly assigned to a global process. However, the product of the two eigenfunctions exhibits significant correlation with the third global eigenfunction.

### Performance evaluation

In order to evaluate the performance of iVAMPnets, we have modified the 10-cube benchmark system to a variable (even) number of subsystems ($N$-cube). We generate trajectories of 100,000 data points for each $N$-cube and train ten instances of both, VAMPnets and iVAMPnets, on it. We recorded the training to be converged when the validation score did not improve by 0.25% compared to the best score so far for 5 training epochs in a row. The results are evaluated first by checking that the global VAMP-E score is indeed converged to its optimum (Fig. 8a), which is true for all instances of trained iVAMPnets. Regular VAMPnets expectedly fail to scale to larger numbers of subsystems – in this case, they do not converge for 8 subsystems and beyond as that corresponds to a number of states larger or equal to $2^8 = 256$. The performance is now evaluated in terms of elapsed real time for training (Fig. 8b). We find that expectedly, both methods have increasing time demands for growing numbers of subsystems or states. We note that the elapsed time for VAMPnets with $\geq 8$ subsystems could not be evaluated due to the failed training procedure. Furthermore, iVAMPnets slightly outperform VAMPnets, which may be caused by the following features of the benchmark system: a) The $N$-cube consists of fully independent subsystems, therefore iVAMPnets can find a domain decomposition quickly. b) The number of states per subsystem in iVAMPnets is just 2, i.e., the neural network parameters can be learned from less data (given a domain decomposition) as compared to a VAMPnet that need to be trained on all transitions between $2^N$ states.

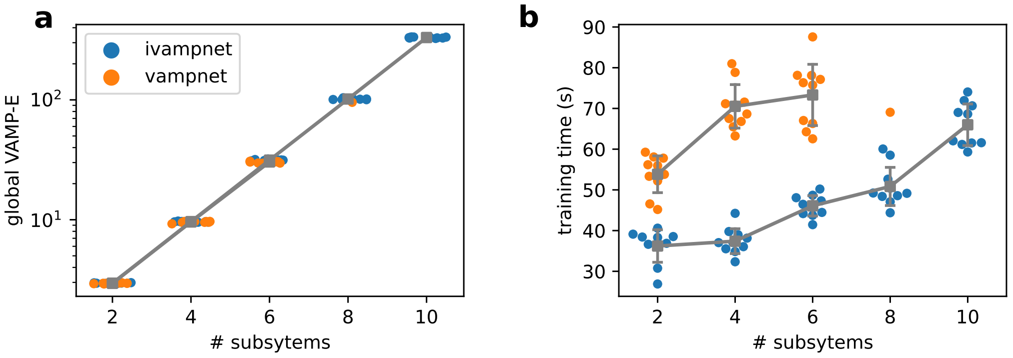


**Figure S8:** Performance evaluation of iVAMPnets compared to VAMPnets using an $N$-cube. **a)** Global VAMP-score as function of the number of subsystems. **b)** Training time (in real time) for both methods, as a function of the number of subsystems. Mean and standard deviations are shown in grey. VAMPnets fail to consistently predict a valid score for 8 subsystems and beyond.

### MD setups

The used MD data sets were generated with the following properties: The synaptotagmin C2A data set [1] consists of 92 trajectories with a length of 2 $\mu$s each, adding up to 184 $\mu$s total simulation time. Simulations were conducted with the CHARMM36 force field [8] in explicit solvent and the NPT ensemble at 300K and 1 bar. Trajectories were seeded from a smaller precursive data set which was based on PDB-ID 2R83 [9]. The villin data set [6] consists of a single trajectory of 125 $\mu$s length that was started from the unfolded structure. Simulations were performed with the CHARMM22* force field [10] in explicit solvent and the NVT ensemble at 360K.
